# NDV-3A vaccination protects mice from *Candida albicans* biofilm infection

**DOI:** 10.1101/471771

**Authors:** Abdullah Alqarihi, Shakti Singh, John E. Edwards, Ashraf S. Ibrahim, Priya Uppuluri

## Abstract

NDV-3A, a novel fungal vaccine undergoing clinical trials, contains a recombinant version of the *Candida albicans* rAls3 N-terminus protein (rAls3p-N) in aluminum hydroxide. In a Phase 1b/2a clinical trial, NDV-3A protected women from recurrent vulvovaginal candidiasis. Here, we reveal that active immunization in mice with NDV-3A induces high titers of anti-rAls3p-N antibodies that interfere with *C. albicans* ability to adhere to and invade endothelial cells, and form biofilm *in vitro*. The anti-rAls3p-N antibodies also protect against *C. albicans* biofilm formation in a central venous catheter mouse model. Finally, anti-rAls3p-N antibodies significantly inhibit yeast dispersal from the hyphal layers of biofilms. Overall these preclinical studies suggest that NDV-3A may serve as an immunotherapeutic strategy for prevention of infections on indwelling medical devices.

## Introduction

*Candida* species are the most common cause of human fungal infections, causing a range of debilitating mucocutaneous infections to life-threatening invasive diseases. Among *Candida* spp., *Candida albicans* is the most common infectious agent, and a frequent cause of fungal genitourinary diseases such as vulvovaginal candidiasis, which is diagnosed in up to 40% of women with vaginal complaints in the primary care settings ^1^. In a hospital setting, hematogenously disseminated candidiasis represents the third-to-fourth most common nosocomial infection worldwide, and *C. albicans* remains the main etiological agent of invasive candidiasis accounting for over half of all cases ^2^. Hematogenously disseminated candidiasis is life threatening, with associated mortality rates between 30 and 60%, despite antifungal use ^3^. The success of *C. albicans* as a pathogen is due to its possessing a number of virulence characteristics, including avid adherence to both abiotic and host cell surfaces ^4^, the capacity to grow tissue-invading filaments ^5^, and the ability to develop biofilms that are resistant to both immune cells and antifungal therapy ^6^. Blocking any of these key virulence traits could serve as the basis of novel therapeutic interventions, with minimal impact on the commensal homeostasis, and reduced selection pressures for emergence of drug resistance ^7^.

Als3p is a vital protein for *C. albicans* virulence. Expressed on the hyphae, Als3p governs all the aforementioned traits (adherence, invasion and biofilms formation) that lead to host pathogenesis ^8^,^9^. Als3p is also required for iron uptake from the host ^10^. Our group has used Als3p for the development of an antiCandida funga vaccinel called NDV-3A ^11^. NDV-3, containing a His-tagged recombinant version of the *C. albicans* Als3 protein N-terminus (rAls3p-N), formulated with alum, prevents both mucosal and hematogenously disseminated candidiasis in mice ^12-15^. NDV-3 was highly immunogenic, well-tolerated and had minimal minor side effects in a Phase I clinical trial ^16^. Furthermore, the latest version of the vaccine, prepared with rAls3p-N without the His-tag, formulated with alum, (NDV-3A), was equally safe and immunogenic, and protective against vaginal infections, in an exploratory 1b/2a clinical trial of patients with a history of recurrent vulvovaginal candidiasis (RVVC) ^11^. Subsequently, our group showed that serum antibodies from patients who responded to NDV-3A (but not those of the unresponsive patients) could prevent *C. albicans* adhesion and biofilm formation on plastic, and invasion of vaginal epithelial cells, *in vitro*. These phenotypic assays served as surrogate markers for efficacy of the NDV-3A vaccine in patients with RVVC ^17^.

The objective of this study was to evaluate and expand the scope of the NDV-3A vaccine against biofilm development *in vivo*, using a mouse model of *C. albicans* biofilm formation. The NDV-3A vaccination induced high titers of anti-rAls3p-N antibodies in mice, which not only prevented adhesion and invasion of human vascular endothelial cells, but also biofilm formation on- and biofilm dispersal from catheter material *in vitro*. Finally, vaccination with NDV-3A also prevented infection of catheters, inserted in the jugular vein of mice. Our studies thus highlight the potential application of NDV-3A against catheter-associated *C. albicans* infections.

## Results and discussion

### NDV-3A vaccination prevented catheter infection in mice

Given the importance of contaminated central venous catheters in hematogenously disseminated candidiasis, we investigated if serum anti-rAls3p-N antibodies could potentially interfere with *C. albicans* attachment and infection of catheters *in vivo*. NDV-3A (300 μg formulated with 200 μg alum), or alum alone (placebo), was administered subcutaneously (s.c.) in mice harboring jugular vein catheters. Vaccination was performed two times (primary + booster dose) at 3 weeks intervals. The booster dose is required in mice because rodents are normally not colonized with *C. albicans*. The mice vaccinated with NDV-3A developed significantly high anti-rAls3p-N antibodies, compared to mice receiving alum alone. Fig 1A depicts the striking difference in antibody levels at 1:300 dilution of sera from antigen-and placebo-vaccinated mice. Specifically, antibody levels in NDV-3A vaccinated mice sera remained higher than alum-vaccinated sera despite dilutions of >1:2000 (**S1A**). This finding is consistent with our recent exploratory Phase 1b/2a study, where a single intramuscular dose of NDV-3A (300 μg Als3p+Alum) resulted in a high anamnestic response characterized by high antibody titers ^11^.

NDV-3A vaccinated and catheterized mice were infected with *C. albicans* intravenously. Catheterized mice vaccinated with alum alone and infected similarly, served as placebo. Three days post infection, NDV-3A vaccination resulted in ~ 1.5 log reduction in catheter and 0.7 log reduction in kidney fungal burden, respectively (Fig 1B), when compared to placebo-vaccinated mice (kidneys p=0.0006 and catheters p=0.04).

**Figure 1.**
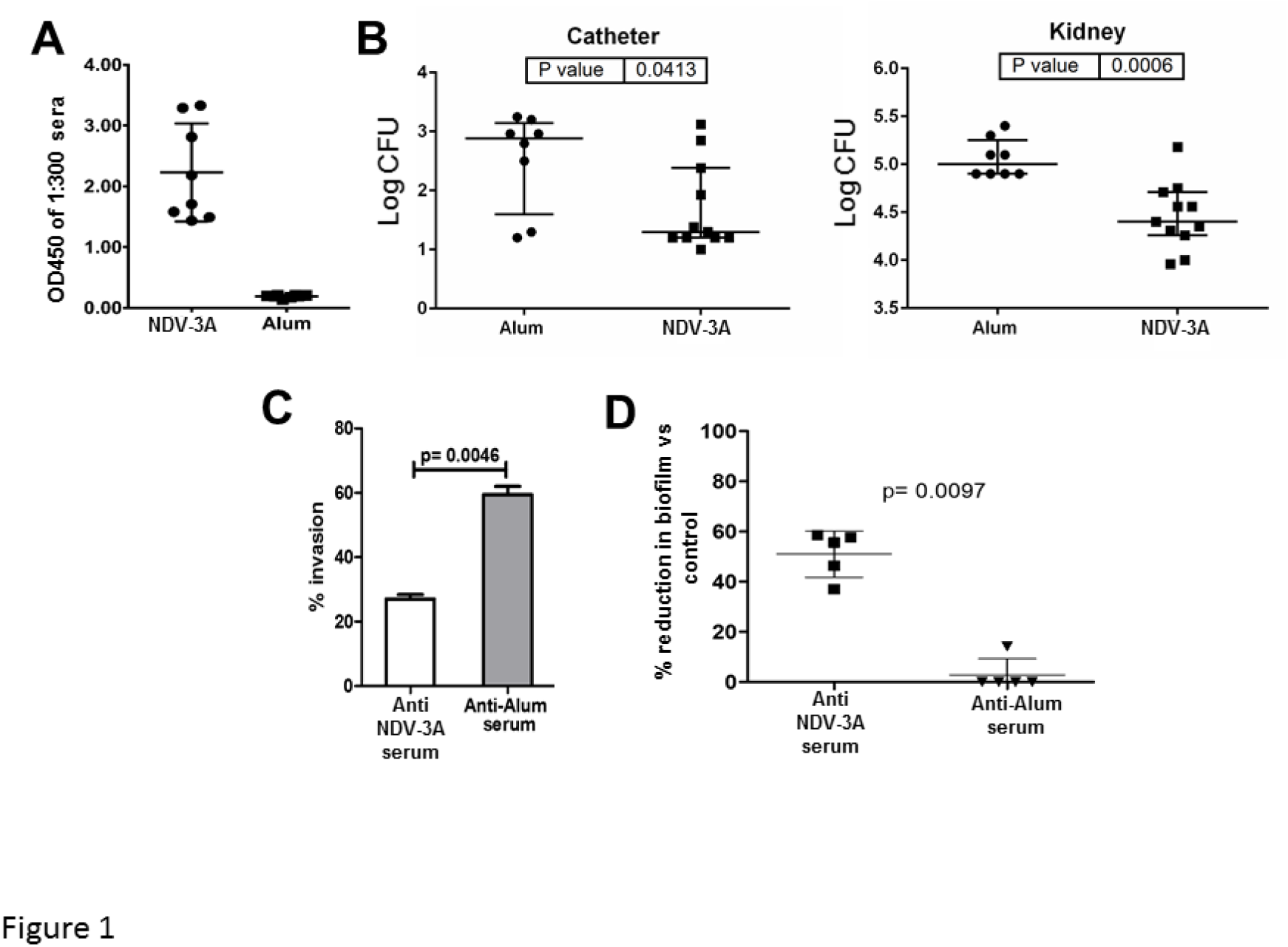
NDV-3A vaccination prevents infection of central venous catheters *in vivo*, and prevents *C. albicans* virulence traits *in vitro*. Vaccination of jugular vein catheterized mice with NDV-3A significantly enhanced anti-rAls3p-N antibodies titers (A). Vaccination protected immunized mice from *C. albicans* infection of jugular vein catheters by ~1.5 log (p=0.04) versus the placebo mice, and significantly (p=0.0006) reduced the fungal burden in mice kidneys after three days, compared to alum-vaccinated mice (B). *C. albicans* pretreated with NDV-3A or placebo antiserum were incubated with HUVEC cells for 90 min, and the number of cells invading the host cells quantified (C). Serum-pretreated *C. albicans* cells were also allowed to develop a biofilm for 24 h, and biofilm metabolic activity quantified by XTT assay (D).

### NDV-3A vaccination prevented *C. albicans* virulence traits, and abrogated biofilm dispersal

Because vaccination yielded high levels of rAls3p-N-specific IgG which were accompanied with decreased *C. albicans* colonization of catheters and reduced kidney fungal burden, we next investigated the role of anti-rAls3p-N antibodies in abrogating *C. albicans* virulence traits presumed to lead to infection in this endovascular catheter model (adhesion and invasion of tissues and/or infection of medical devices such as catheters). Specifically, we investigated the extent of fungal adhesion and invasion of human umbilical vein endothelial cells (HUVEC) since these cells are the first line of defense that *C. albicans* encounters during hematogenous dissemination, in the presence or absence of anti-rAls3p-N sera. Post-vaccination sera from mice which received NDV-3A significantly (p=0.004) reduced invasion of *C. albicans* to HUVEC, by 50%, compared to alum vaccinated sera (Fig 1C). Next, we examined if anti-rAls3p-N antibodies could block biofilm growth on silicone elastomer catheter material. We showed that NDV-3A vaccinated serum abrogated at least 50% of biofilm growth, when added at the time of biofilm initiation, compared to when biofilms were developed in the presence of alum-vaccinated serum (Fig 1D). The sera however failed to kill or disrupt preformed biofilms (data not shown). *C. albicans* biofilms form a robust mesh of yeast and hyphal cells encased in an extracellular matrix, inert to almost all classes of antifungal drugs as well as resistant to immune cell attack ^6^. Our finding that anti-rAls3p-N antibodies prevented adhesion, invasion and biofilm formation is not without premise. First, the sera were heat treated to rule out a role for complement in the experimental outcome. Second, we previously demonstrated that NDV-3A protects mice from VVC by a mechanism that involves the priming of both humoral and adaptive immune responses ^12^. The importance of antibodies targeting virulence traits of *C. albicans* was also demonstrated in animal models against other important antigens including Hyr1p ^18,19^, Sap2 ^20^, and *Candida* cell wall glycopeptides ^21^,^22^. Our results show that murine (this study) and human anti-rAls3p-N antibodies display enhanced efficacy at neutralizing *C. albicans* Als3-associated primary functions of adhesion, invasion and biofilm formation.

As a part of normal biological growth, *C. albicans* hyphae produce budding yeast cells from their lateral septal regions. When released from the biofilm-hyphal mesh, these cells from the lateral septal regions are called biofilm-dispersed cells. Biofilm dispersed cells from biofilm covered catheters have direct access to the blood stream, and are implicated in perpetuating dissemination ^23,24^. Although the antibodies induced by NDV-3A were inactive with fully formed biofilm, incubation with NDV-3A derived antisera prevented formation of lateral yeast cells, thus reducing dispersal of yeast cells from the biofilms, compared to both biofilms treated with sera from alum-vaccinated mice, and commercially available sera (Fig 2A). Specifically, the serum from mice immunized with NDV-3A reduced lateral yeast cell dispersal by 35% (Fig 2B, p=0.001). Overall, these results highlight the potential of NDV-3A vaccination to not only prevent disseminated infections, but also to block the source of dissemination, since biofilms act as a reservoir of dispersing cells directly into the blood stream ^24^. This finding also suggests a role of Als3p, an abundant hyphal cell-surface protein, in lateral yeast cell growth from hyphae.

**Figure 2.**
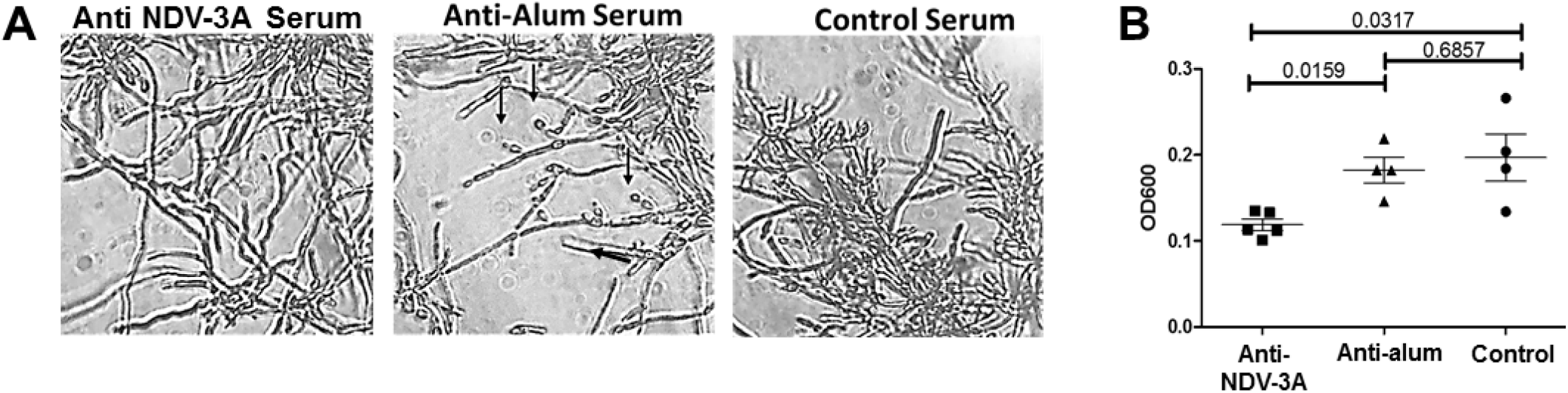
Anti NDV-3A sera prevents biofilm dispersal *in vitro*. Biofilms (6 h) were incubated with the respective (5%) sera for 12 h at 37°C, and aliquots of biofilms visualized microscopically to gauge the effect of anti-NDV-3A serum in preventing biofilm dispersal (A). Turbidity of the media over the biofilms (that represented dispersed cells) were also quantified by measuring OD600 (B).

Collectively, these *in vitro* and *in vivo* studies serve as evidence for the role of antibodies in abrogating disseminated candidiasis. The NDV-3A vaccination induced significant abrogation of *C. albicans* pathogenic traits and reduced subsequent lethality. While this inhibition of virulence traits or protection from candidiasis was not 100%, we submit that this vaccine is not expected to be used as an isolated therapeutic option alone. We postulate that the preventative activity exhibited by vaccination can be further augmented in the presence of antifungal drugs. It is possible that such a combination of the vaccine with antifungal drugs would allow the usage of smaller quantities of antifungal drugs, thereby inhibiting the evolution of antifungal drug resistance. This hypothesis is the subject of future studies by our group.

In summary, our study illuminates the broad therapeutic relevance of anti-rAls3p-N antibodies generated after vaccination, in preventing *C. albicans* virulence both *in vitro* and *in vivo*, including device-associated candidiasis. These studies also include efforts to directly target a specific virulence determinant influencing host-pathogen interactions. We hope to provide additional concepts for further developing vaccination strategies for fungal pathogens in general. Currently, no vaccines have been approved for human fungal pathogens.

## Material and methods

### *Candida* Strain and NDV-3A

*C. albicans* SC5314 was routinely cultured overnight in yeast peptone dextrose (YPD) broth (Difco) at 30°C with shaking, prior to use for *in vitro* assays. In all studies, *C. albicans* cells were washed twice with endotoxin-free Dulbecco’s PBS, suspended in PBS or yeast nitrogen base (YNB, Difco), and counted with a hemocytometer to prepare the final inoculum. *Candida albicans* SC5314 was the source of the N-terminus of Als3 used to develop the NDV-3A vaccine^11^. NDV-3A was kindly provided by NovaDigm Inc., where the vaccine was generated.

### Mice immunizations and infection models

The NDV-3A vaccine was formulated by mixing 300 μg of *Candida albicans* rAls3p-N with alum (200 μg) adjuvant. The recombinant Als3 protein was produced in *Saccharomyces cerevisiae*, as described previously. We used 4-6 weeks jugular vein catheterized male C57BL/6 mice (Charles River labs, Wilmington, MA) in this study. For mice immunization, NDV-3A vaccine or alum was injected subcutaneously (s.c.) into mice at day 0 and boosted with a similar dose on day 21. For serum collection, mice were bled 14 days following the boost. The immunized mice were infected via tail vein with 5×10^5^ cells of *C. albicans* 15 days after the boost. Three days post infection, mice were sacrificed and the catheters within the jugular vein and the surrounding tissue as well as kidneys were recovered. Catheters along with the surrounding tissues were weighed, cut into small pieces, and homogenized, as were the kidneys. The kidney homogenates were diluted and quantitatively cultured overnight for measurement of CFU. All the procedure conducted on mice were approved by IACUC of Los Angeles Biomedical Research Institute at Harbor-UCLA Medical Center (Protocol number 11672). All experiments were performed in accordance with relevant guidelines and regulations.

### Measurement of antibody titers

The anti-rAls3p-N antibody titer was measured by performing a standard ELISA test ^25^. Briefly, the wells of a 96-well microtiter plate was coated with rAls3p-N (5 μg/ml) as a bait, washed with phosphate-buffered saline (PBS) containing 0.05% Tween 20 (PBST) and blocked by incubation with 200 μl of 1% bovine serum albumin (BSA) in PBST. After additional washes, various dilutions of mice sera (NDV-3A vaccinated or placebo) were allowed to interact with the protein for 2 hr at 37°C. After three washes, plates were incubated at 37°C for 2 h with peroxidase-conjugated anti-human IgG (Sigma Chemical Co., St. Louis, Mo.) diluted 1:1,000 in PBST. Finally, after three washes, freshly prepared solution of the substrate 1 mg/ml PNPP (p-nitrophenyl phosphate) was added, and plates incubated for 30 min – 2 hr in dark. The reaction was measured at OD405, using a spectrophotometer. Differences in the ELISA readings between the two vaccinated mice groups were plotted in Microsoft Excel.

### HUVEC Invasion assays

HUVEC were obtained by a modification of the method of Jaffe et al. ^26^. The cells were grown in M-199 medium with 2 mM L-glutamine, penicillin, and streptomycin (Gibco) and supplemented with 10% fetal bovine serum and 10% bovine calf serum (Gemini BioProducts, Woodland, Calif.). Second-or third-passage cells were grown to confluence on 0.2% gelatin matrix-coated circular coverslips, in 24-well tissue culture plates. All incubations were in 5% CO_2_ at 37°C. On the day of the assay, 1 ml of 1×s10^5^ *C. albicans* cells in the tissue culture medium, were pretreated with sera (from NDV-3A-vaccinated or placebo mice or control commercial serum) at a final concentration of 5%. To measure for the role of compliment on *C. albicans* of HUVEC, aliquots of mice sera were independently heated in a heat block set to 55°C for 1 h, and incubated with *C. albicans* as above. After 30 min of incubation at 37°C to allow for germination and expression of Als3p, the *C. albicans-sera* mixture was added to the wells containing the endothelial cells (after aspirating the medium over the host cells), and incubated for 90 min at 37°C in 5% CO_2_. The plates were washed twice with sterile PBS, and the coverslips stained with 25 μg/ml Concanavalin A (ConA) for 30 min at 37°C. The number of *C. albicans* cells adhered to the endothelial monolayer was counted under an inverted light microscope. The extent of endothelia cell invasion was visualized by differential staining and counting: absence of stain in fungi invading the mammalian cells versus stained, non-invading fungal cells ^17^. The differential counting was done under a Confocal Scanning Laser Microscope (Leica SP2) by overlaying the brightfield image with a 594 nm excitation filter (red laser) for ConA. At least 50 fields were randomly counted for each experimental condition. Extent of invasion by anti-rAls3p-N sera and anti-alum sera were plotted as percent invasion compared to IgG isotype control sera.

### Biofilm-related assays

Biofilms were developed on silicone elastomer (SE) pieces, in the presence/absence of serum, with modifications ^23^. Briefly, circular SE pieces with the approximate diameter of a 96-well were prepared by punching the SE sheets with a paper puncher. These autoclaved and sterile SE circles were pre-incubated with 100% fetal bovine serum overnight at room temperature, washed twice with sterile water, and then introduced into the wells of a 96-well plate, where they fit snugly. *C. albicans* blastospores (2 × 10^5^ cells/ml in YNB medium) were added to wells containing 5 μL of respective NDV-3A, alum vaccinated mice sera or commercial serum (each 5% serum vol/vol), and incubated at 37°C. Biofilm was then allowed to develop in the SE-bottomed wells for 24 h at 37°C, washed once with sterile water, and estimated using the XTT calorimetric assay, that measures the metabolic activity of the cells^27^. Data was plotted as percent reduction in biofilm growth versus the control.

For measuring the impact of anti-rAls3p-N antisera on biofilm dispersal, *C. albicans* biofilms were developed for 6 h in 96-well microtiter plates, after which the wells were washed twice with sterile water. YNB medium with 5% sera from NDV-3A or alum-vaccinated mice, and commercially available serum (as control) was added to the wells, and the biofilms allowed to grow for another 12 h at 37°C. Media over the biofilms were aspirated and turbidity (OD600) quantified. At the same time, aliquots of biofilms were placed on a glass slide to visualize under a bright field microscope at 40X magnification, and images captured.

### Statistical Analysis

All *in vitro* studies were performed in triplicate at a minimum, with two biological replicates. Different groups were compared using the non-parametric Mann–Whitney test for comparison of unmatched groups. Data were analyzed in GraphPad Prism software (La Jolla, CA, USA), and a *p*-value <0.05 was considered statistically significant.

For *in vivo* studies, a power analysis for a two-tailed students t-test was used which results in a 90% power to detect a log difference in tissue and catheter fungal burden.

## Acknowledgments

We would like to thank John P. Hennessey for kindly providing us with NDV-3A vaccine. This work was supported by NIH grant R01 AI063382 to JEE, and a UCLA CTSI Institutional KL2 Translational Science Award to PU. The funders had no role in study design, data collection and analysis, decision to publish, or preparation of the manuscript.

## Author contributions

AA performed the in vitro experiments, SS assisted the in vivo studies, JEE provided valuable scientific input, ASI helped in the design of the project and write the manuscript, PU designed and performed the studies and wrote the manuscript.

## Conflict of Interest

JEE and ASI are founders and shareholders of NovaDigm Therapeutics Inc. a company that is developing the NDV-3A as a fungal vaccine. All other co-authors have no formal association with NovaDigm.

**S1. Anti-NDV-3A vaccinated mice sera (n= 8 per group) displayed significantly high anti-rAls3p-N antibod**y **titers compared to alum treated mice, even at dilutions as high as 1:2000**.

